# The genetic architecture of protein stability

**DOI:** 10.1101/2023.10.27.564339

**Authors:** Andre J. Faure, Aina Martí-Aranda, Cristina Hidalgo-Carcedo, Jörn M. Schmiedel, Ben Lehner

## Abstract

There are more ways to synthesize a 100 amino acid protein (20^100^) than atoms in the universe. Only a miniscule fraction of such a vast sequence space can ever be experimentally or computationally surveyed. Deep neural networks are increasingly being used to navigate high-dimensional sequence spaces. However, these models are extremely complicated and provide little insight into the fundamental genetic architecture of proteins. Here, by experimentally exploring sequence spaces >10^10^, we show that the genetic architecture of at least some proteins is remarkably simple, allowing accurate genetic prediction in high-dimensional sequence spaces with fully interpretable biophysical models. These models capture the non-linear relationships between free energies and phenotypes but otherwise consist of additive free energy changes with a small contribution from pairwise energetic couplings. These energetic couplings are sparse and caused by structural contacts and backbone propagations. Our results suggest that artificial intelligence models may be vastly more complicated than the proteins that they are modeling and that protein genetics is actually both simple and intelligible.

## Introduction

Massively parallel experiments allow the effects of single amino acid (aa) changes in proteins to be comprehensively quantified ^1,2^. Similarly, experimental analysis of double mutants is feasible, at least for small proteins ^3,4^. The analysis of higher order mutants, however, quickly becomes infeasible due to the combinatorial explosion of possible genotypes. For example, the number of ways to combine one mutation at 34 different sites in a protein is 2^34^ ∼ 1.7 x 10^10^. Experimental exploration of such a large number of genotypes is extremely challenging given current technology, which to-date has experimentally analyzed sequence spaces up to ∼ 10^6^ ^3,5^.

Moreover, combining random mutations in even moderate numbers nearly always results in unfolded proteins ^6,7^. For example, only 2-8% of 5 aa variants and <0.2% of 10 aa variants in a small protein domain are expected to be folded (n = 2 domains, Fig. 1a, Extended Data Fig. 1a). Sampling even tens of millions of random combinatorial genotypes in most proteins will therefore provide almost no information about genetic architecture and will not be useful for training and evaluating predictive models beyond testing the trivial prediction that most genotypes are unfolded.

**Figure 1.**
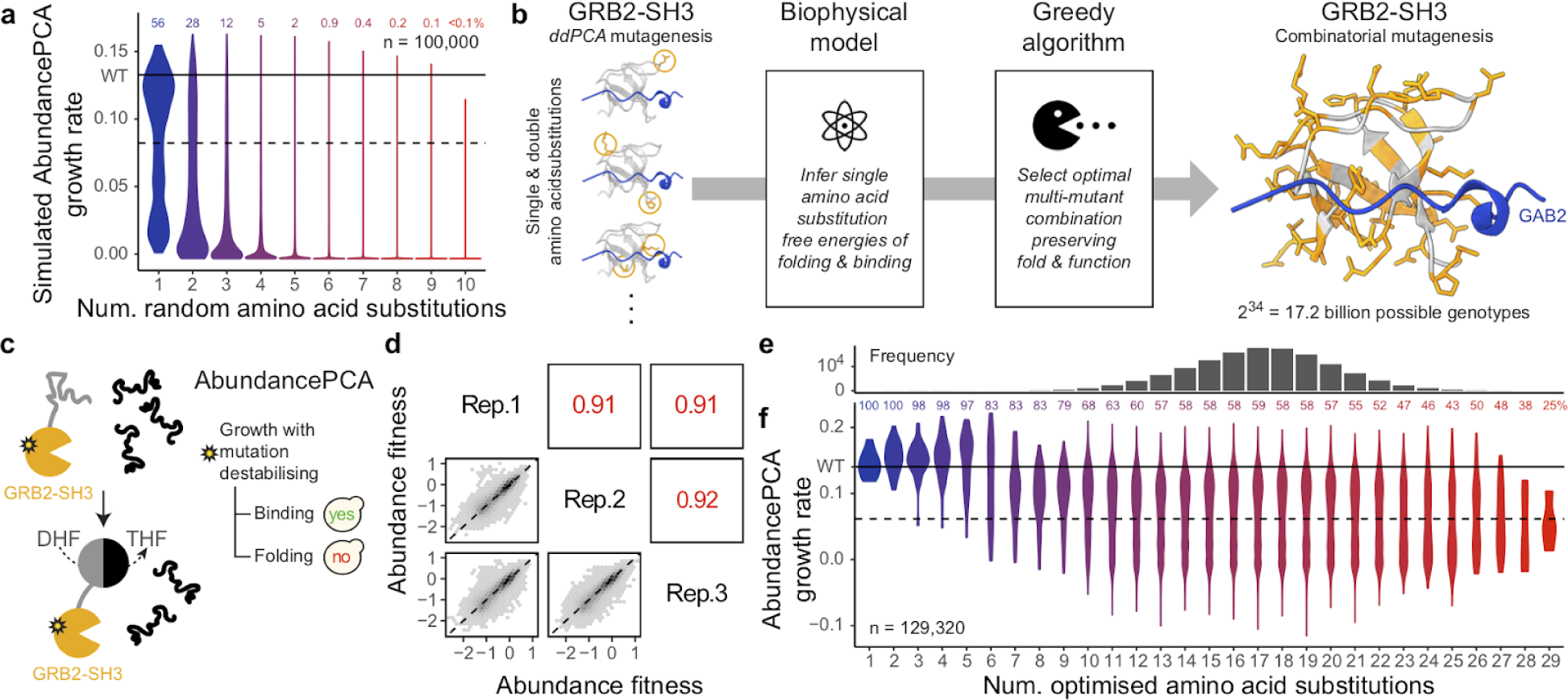
An efficient strategy to explore high-dimensional protein sequence space and enrich multi-mutants for conserved fold and function. **a,** Violin plot showing distributions of simulated AbundancePCA growth rates (assuming additivity of individual inferred folding free energy changes ^23^) versus number of random amino acid substitutions (n = 100,000). Violins are scaled to have the same maximum width. b, DMS data, biophysical model and algorithm used to select a set of single aa substitutions for combinatorial mutagenesis. A shallow double mutant library of GRB2-SH3 protein variants was assayed by AbundancePCA (see panel c) and BindingPCA (see Fig. 5b; in combination referred to as ddPCA) followed by biophysical modeling to infer single aa substitution free energy changes of folding and binding ^23^. We used this model together with a greedy algorithm to select a set of 34 single aa substitutions which, when combined, would simultaneously maximize both the predicted AbundancePCA and BindingPCA growth rates i.e. preserving both fold and function. 3D structure of GRB-SH3 (PDB: 2VWF) indicating the 34 combinatorially mutated residues (orange) and GAB2 ligand (blue) is shown at right. c, Overview of AbundancePCA on protein of interest (GRB2-SH3) ^23^. yes, yeast growth; no, yeast growth defect; DHF, dihydrofolate; THF, tetrahydrofolate. d, Scatter plots showing the reproducibility of fitness estimates from AbundancePCA. Pearson’s *r* indicated in red. Rep., replicate. e, Histogram showing the number of observed aa variants at increasing Hamming distances from the wild type where the x-axis is shared with panel f. f, Violin plot showing distributions of AbundancePCA growth rates inferred from deep sequencing data versus number of amino acid substitutions. In panels a and f, the percentage of folded protein variants (predicted fraction folded molecules > 0.5) is shown at each Hamming distance from the wild type.

One strategy for exploring high-dimensional sequence spaces is to use generative artificial intelligence. Deep neural networks with millions of fitted parameters have proven successful for diverse prediction and protein design tasks, including predicting the effects of combinatorial mutants ^8–18^. However, these models have extremely complicated and difficult to interpret architectures and alone provide little insight into the actual genetic architecture of proteins.

It could be that the extreme complexity of these models reflects an underlying extreme complexity of protein genotype-phenotype landscapes. Alternatively, the actual genetic architecture of proteins might be much simpler than is evidenced by these models. Such simplicity is suggested by the success of quite simple statistical models in four dimensional sequence spaces ^19^ and the use of Potts Hamiltonian models that only consider positional conservation and pairwise co-variation in multiple sequence alignments of protein families to predict protein structures ^20,21^ and activity ^22^.

Here, we use an experimental design that enriches functional protein sequences to explore the fundamental genetic architecture of high-dimensional protein sequence spaces with >30 dimensions and >10^10^ genotypes. Despite the complexity of deep learning models we find that protein architectures are remarkably simple, with additive biophysical models providing very good predictive performance. Quantifying the pairwise energetic couplings between mutations further increases predictive power, providing excellent performance in high-dimensional genotype spaces. These couplings are sparse and related to protein 3D structures. The genetic architecture of at least some proteins is therefore very simple, with additive energetics and a minor contribution from sparse pairwise structural couplings.

## Results

### Experimentally sampling a 10^10^ sequence space

We previously showed that the energetic effects of thousands of individual mutations on the stability of a protein can be measured *en masse* using pooled variant synthesis, selection and sequencing experiments ^23^. In these experiments, the effect of each mutation on the cellular abundance of a protein is measured in the wild-type (WT) protein and also in a small number of variants with different fold stabilities. For example, using a shallow double mutant library we could infer the changes in Gibbs free energy of folding (ΔΔ*G*_f_) for nearly all mutations (1056/1064 = 99%) in the C-terminal SH3 domain of the adaptor protein GRB2 ^23^. Similar massively parallel measurements of single mutant fold stabilities have now been made for other signaling domains including the oncoprotein KRAS ^23,24^ and, *in vitro,* for many, mostly prokaryotic, small domains ^5^.

Combining random mutations in the GRB2-SH3 domain very quickly results in unfolded proteins, with ∼98% and >99.9% of genotypes with 5 and 10 mutations expected to be unfolded (based on additive energies, see Fig. 1a). This rapid decay of stability as mutations are combined is consistent with experimental measurements of other proteins ^6,7^. To experimentally explore folded genotypes in high-dimensional sequence spaces we therefore used a heuristic technique to enrich for conserved fold and function in combinatorial variants. For each possible starting single aa substitution, we iteratively selected additional substitutions – one per residue position – that simultaneously maximizes the resulting combinatorial mutant’s predicted abundance and binding to an interaction partner (see Methods). For GRB2-SH3, the largest set of mutations predicted to preserve both molecular phenotypes consisted of 34 single aa substitutions: 25 in surface residues (relative solvent-accessible surface area (RSASA) ≥ 0.25), three in the protein core (RSASA < 0.25), and six mutations in the GAB2 ligand binding interface (ligand distance < 5 Å, Fig. 1b, right).

We synthesized a library containing all combinations of these 34 mutants and quantified the cellular abundance of a small sample of the 2^34^ ∼ 1.7 x 10^10^ genotypes using a highly validated pooled selection, abundance protein fragment complementation assay (AbundancePCA, aPCA ^23–25^). In total, we obtained triplicate abundance measurements for 129,320 variants, which is 0.0007% of the sequence space. The measurements were highly reproducible (Pearson’s *r* > 0.91, Fig. 1d, see Methods).

The symmetrical pod-like shape of the genotype frequency landscape, with the number of genotypes peaking at the intermediate Hamming distance of 17 i.e. equidistant from the WT (0th order) and 34th order mutant – is recapitulated in the bottlenecked library results (Fig. 1e). Median abundance measurements decrease with increasing number of amino acid (aa) substitutions, but there are still thousands of genotypes with many mutations that nevertheless maintain abundance scores that are indistinguishable from that of the WT protein (*n* = 2,706 with >20 mutations, nominal *P* > 0.05, Fig. 1f).

### Accurate combinatorial genetic prediction with simple biophysical energy models

Quantifying the effects of a large number of multi-mutants allowed us to test the generalisability of genotype-phenotype models, including in regions of the genetic landscape far beyond the local neighborhood used for training. For model building and evaluation we restricted all analyses to variants with quantitative measurements in all three biological replicates (*n* = 71,233). Surprisingly, our original biophysical model (Fig. 2a) trained on abundance and ligand binding selections (doubledeepPCA, ddPCA) quantifying the effects of single and double aa mutants only ^23^ explains as much as half of the fitness variance in combinatorial multi-mutants (*R*^2^ = 0.5, Fig. 2b, lower right panel), where the vast majority (94%) include at least 13 aa substitutions in the wild-type sequence. The only trained parameters in this simple model are Gibbs free energy terms for the wild type (Δ*G*_f_) and single aa substitutions (ΔΔ*G*_f_), and a two-parameter (affine) transformation relating the fraction of folded molecules to the AbundancePCA score (fitness; Fig. 2a). That such a large proportion of phenotypic variance is explained by an additive biophysical model (no specific epistasis/interactions) that is trained on genotypes that only contain 1 or 2 genetic changes suggests that the energetic effects of mutations in proteins are largely context-independent.

**Figure 2.**
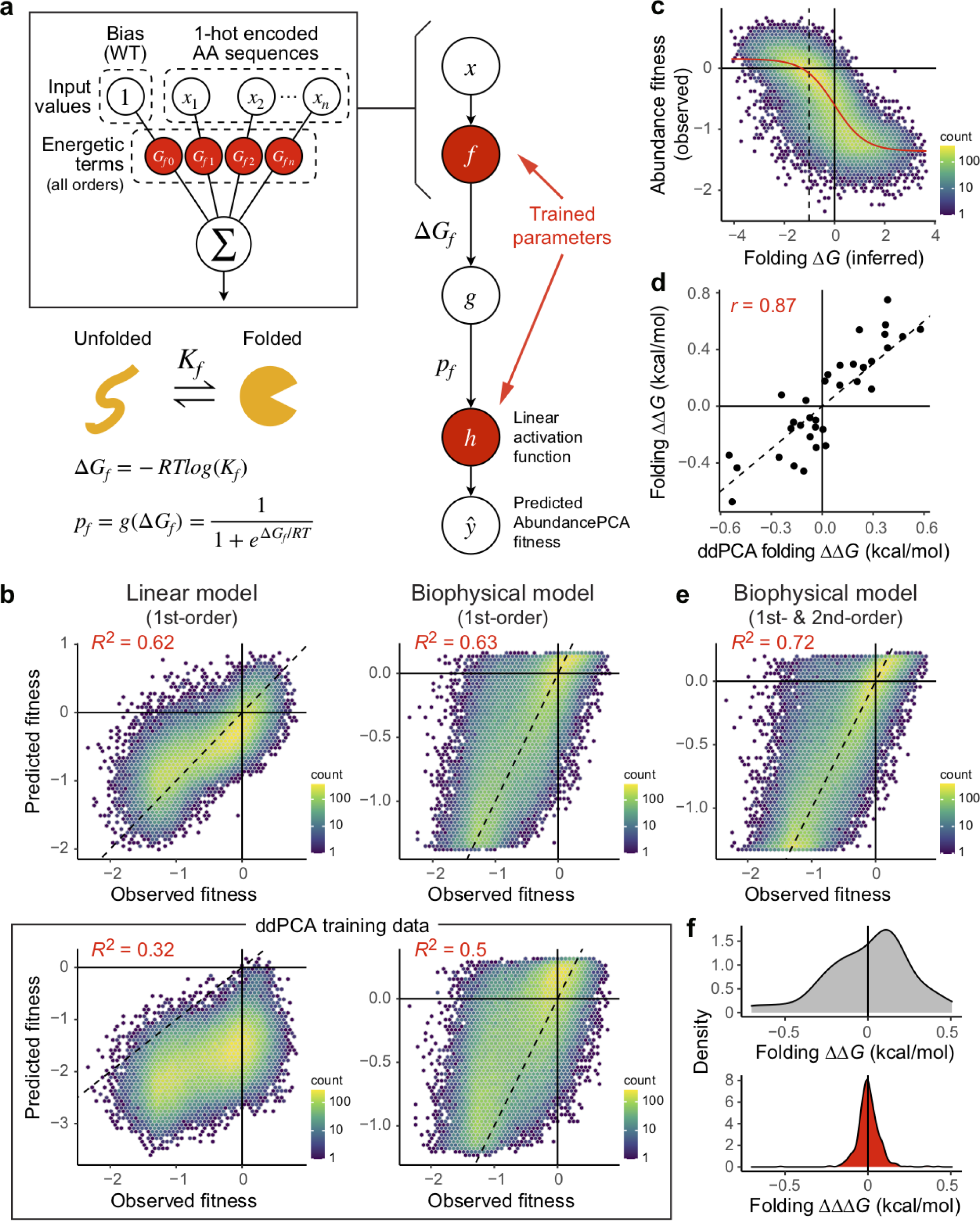
Biophysical modeling of protein abundance to infer folding free energy changes and energetic couplings. **a**, Two-state equilibrium and corresponding neural network architecture used to fit thermodynamic model to AbundancePCA data (bottom, target and output data), thereby inferring the causal changes in free energy of folding associated with single amino acid substitutions (top, input values). Δ*G*_f_, Gibbs free energy of folding; *K*_f_, folding equilibrium constant; *p*_f_, fraction folded; *g*, non-linear function of Δ*G*_f_; R, gas constant; *T*, temperature in Kelvin. **b**, Performance of first-order linear models (left column) and first-order biophysical models (right column) evaluated on GRB2-SH3 combinatorial AbundancePCA data. The upper row indicates results of models that were trained on a subset of the same combinatorial DMS data. The lower row indicates results of models that were trained on GRB2-SH3 ddPCA data consisting of single and double aa substitutions only ^23^. *R*^2^ is the proportion of variance explained. **c**, Non-linear relationship (global epistasis) between observed AbundancePCA fitness and changes in free energy of folding. Thermodynamic model fit shown in red. **d**, Comparisons of the model-inferred free energy changes to previously reported estimates using GRB2-SH3 ddPCA data ^23^. Pearson’s *r* is shown. **e**, Performance of biophysical model including all first- and second-order genetic interaction (energetic coupling) terms/coefficients. See Extended Data Fig. 2 for plots of the residuals versus fitted values for linear and biophysical models of first- and secondorder. **f**, Distributions of folding free energy changes (ΔΔ*G*, gray) and pairwise energetic couplings (ΔΔΔ*G*, red).

On the other hand, a linear model – that implicitly assumes mutation effects combine additively at the phenotypic level in multi-mutants – trained on the same ddPCA data – performs significantly worse (*R*^2^ = 0.32). The linear model also systematically underestimates the observed phenotypic effects of mutant combinations (Fig. 2b, lower left panel), a consequence of not accounting for the scaling of mutational effects due to protein thermodynamics (global epistasis ^26^). For example, introducing a destabilizing mutation in an already-unfolded protein has no effect on the fraction of folded molecules (lower plateau of model in Fig. 2c), which is not captured by a linear model. These results demonstrate a key advantage of fitting biophysical models: accounting for global epistasis improves the generalisability of predictions beyond the (local neighborhood) of the training data.

Fitting linear and biophysical models to the combinatorial data improves performance by 30% and 13% respectively (Fig. 2b, upper panels), likely due to the greater amount of training data and (relevant) genetic backgrounds in which the effects of each single aa are quantified: ∼50% of all variants in the library (*n* ∼ 30,000) include – and therefore report on the effects of – any given one of the 34 single aa substitutions i.e. almost three orders of magnitude more measurements per single mutant compared to the relatively shallow (ddPCA) library. Although the fraction of variance explained by first-order linear and biophysical models is comparable (*R*^2^ = 0.62 and 0.63, Fig. 2b, upper panels), the biased regression residuals in the case of the linear model show this model is less appropriate (Extended Data Fig. 2a). The biophysical model provides an excellent fit to the data, faithfully capturing the global non-linear relationship (global epistasis) between observed AbundancePCA fitness and inferred changes in free energy of folding (Δ*G*_f_, Fig. 2c). There is also very good agreement between free energy changes (model parameters) inferred from combinatorial and ddPCA datasets (Pearson’s *r* = 0.87), but the former tend to be more extreme, once again demonstrating the utility of assaying the effects of mutations in greater numbers of genetic backgrounds thereby allowing their energetic effects to be more accurately estimated (Fig. 2d).

### Pairwise energetic couplings improve genetic prediction

We next tested whether quantifying non-additive energetic couplings between mutations improved predictive performance. In our combinatorial dataset each pair of mutations is present in a median of 17,923 genotypes, allowing robust measurement of second-order genetic interaction terms (energetic couplings, ΔΔΔ*G*_f_ ^27,28^). Including all second-order energetic couplings improves model performance by an extra 9% (*R*^2^ = 0.72), consistent with expectations that pairwise effects are an important source of specific epistasis in proteins ^28^ (Fig. 2e). Whereas first-order terms are stronger in magnitude and biased towards destabilizing effects, second-order energetic couplings tend to have milder effects centered on zero (Fig. 2f).

### Strong energetic couplings are due to physical contacts and backbone propagations

Measured at the phenotypic level, genetic interactions in proteins have previously been shown to reflect – at least in part – protein structures ^3,29–31^. Combining combinatorial deep mutational scanning with biophysical modeling allowed us to infer a total 561 pairwise energetic couplings, providing an opportunity to interrogate their mechanistic origins and relationship to protein structure. Comparing coupling energy magnitude (absolute folding ΔΔΔ*G*_f_) to the 3D distance separating mutation pairs in the folded structure (minimal side chain heavy atom distance) reveals an L-shaped distribution with the strongest energetic couplings occurring between structurally proximal residues (Fig. 3a; see also Extended Data Fig. 3a). The top five energetic couplings all involve pairs of residues within 5.5 Å and 15/20 (75%) of the top 20 energetic couplings involve pairs separated by less than 8 Å. Although there is a weak anti-correlation between contact distance and coupling energy strength (Spearman’s *ρ* = -0.12, Fig. 3a), this trend breaks down for pairs that are not proximal in the primary sequence (Spearman’s *ρ* = -0.02, backbone distance >5 residues). In other words, disregarding residue pairs that are nearby in the (1D) aa sequence, separation distance in 3D space is not predictive of coupling strength.

**Figure 3.**
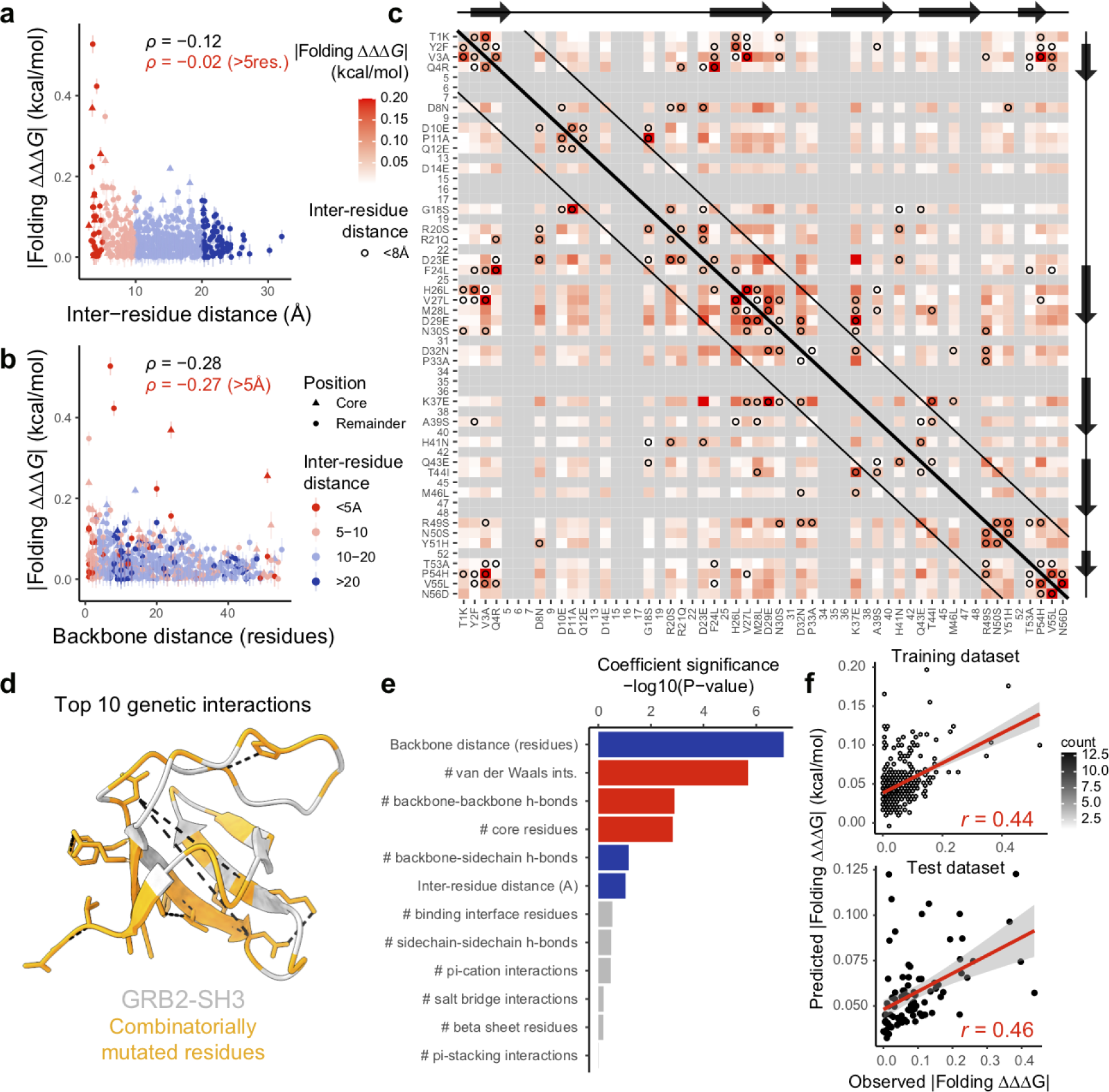
Structural determinants of energetic couplings inferred from combinatorial mutagenesis. **a**, Relationship between folding coupling energy strength and minimal inter-residue side chain heavy atom distance. Error bars indicate 95% confidence intervals from a Monte Carlo simulation approach (*n* = 10 experiments). Points are coloured by binned inter-residue distances (see legend in panel b). Spearman’s *ρ* is shown for all couplings as well as those involving pairs of residues separated by more than five residues in the primary sequence (red). Core residues are indicated as triangles. **b**, Relationship between folding coupling energy strength and linear sequence (backbone) distance in number of residues. **c**, Interaction matrix indicating folding coupling strength as well as pairwise structural contacts in GRB2-SH3 (PDB: 2VWF, minimal side chain heavy atom distance < 8 Å, black circles). Gray cells indicate missing values (non-mutated residues) and constitutive single aa substitutions are indicated in x- and y-axis labels. Black diagonal lines demarcate residue pairs that are distal in the primary sequence (backbone distance > 5 residues). Reference secondary structure elements (arrow, β-strand) are shown at top and right. **d**, 3D structure of GRB-SH3 (PDB: 2VWF) indicating the top 10 energetic couplings with black dashed lines. Combinatorially mutated residues are shown in orange. **e**, Bar plot showing ranked features from linear model to predict folding coupling strength. Bar width indicates coefficient significance. Blue, positive coefficient; red, negative coefficient; gray non-significant (nominal *P* > 0.05). **f**, Correlation between linear model predicted and observed folding coupling strength for training (top) and test data (bottom). Pearson’s *r* is shown.

On the other hand, comparing coupling strength to separation distance between residues in the primary sequence (along the peptide backbone) reveals a significant inverse relationship that extends over very large distances (Spearman’s *ρ* = -0.28) and is robust to the exclusion of direct physical contacts between residues (< 5 Å, Spearman’s *ρ* = -0.27, Fig. 3b; see also Extended Data Fig. 3b). The observation that mutation effects can be modulated by the sequence context at distal sites and that the strength of this coupling depends in part on the number of intervening residues – but not 3D distance – suggests that energetic perturbations are particularly efficiently transmitted over long distances along the covalent bond structure of the peptide backbone.

The interaction matrix in Figure 3c summarizes these observations: the strongest energetic couplings coincide with direct physical contacts (black circles, see also Fig. 3d), and energetic coupling strength decays along the protein backbone (Fig. 3c, near-diagonal vs. far off-diagonal cells). The matrix also highlights physical interactions between secondary-structural elements as hotspots for strong energetic couplings.

In order to disentangle the relative importance of these different potential structural determinants of energetic coupling strength, we gathered a collection of quantitative features describing both the number and type of chemical bonds or interactions existing between the atoms of pairs of residues as well as their relative positions in the folded structure (Fig. 3e). A linear regression model based on these 12 structural features is predictive of coupling strength (Fig. 3f, see Methods). Importantly, the same model performs similarly well when evaluated on a held out, non-overlapping set of inferred energetic couplings derived from an independent combinatorial mutagenesis experiment (Pearson’s *r* = 0.46, *R*^2^ = 0.21). This suggests that, despite its simplicity, the integrated model captures structural determinants of energetic coupling strength and that energetic couplings are caused by structural interactions.

### Energetic couplings decay along the protein backbone

To directly test the hypothesis that inter-residue backbone distance is associated with energetic coupling strength independently of 3D contact distance, we designed a combinatorial saturation mutagenesis library involving all possible mutations at four physically proximal surface residues in the same secondary structure element (Fig. 4a, Extended Data Fig. 4, see Methods). We reasoned that energetic couplings due to the propagation of perturbations along the protein backbone should also be apparent among solvent-facing residues. In total, we obtained abundance measurements for 138,157 variants (86% of the sequence landscape) and the measurements were highly reproducible (Pearson’s *r* > 0.89, Fig. 4b, see Methods).

**Figure 4.**
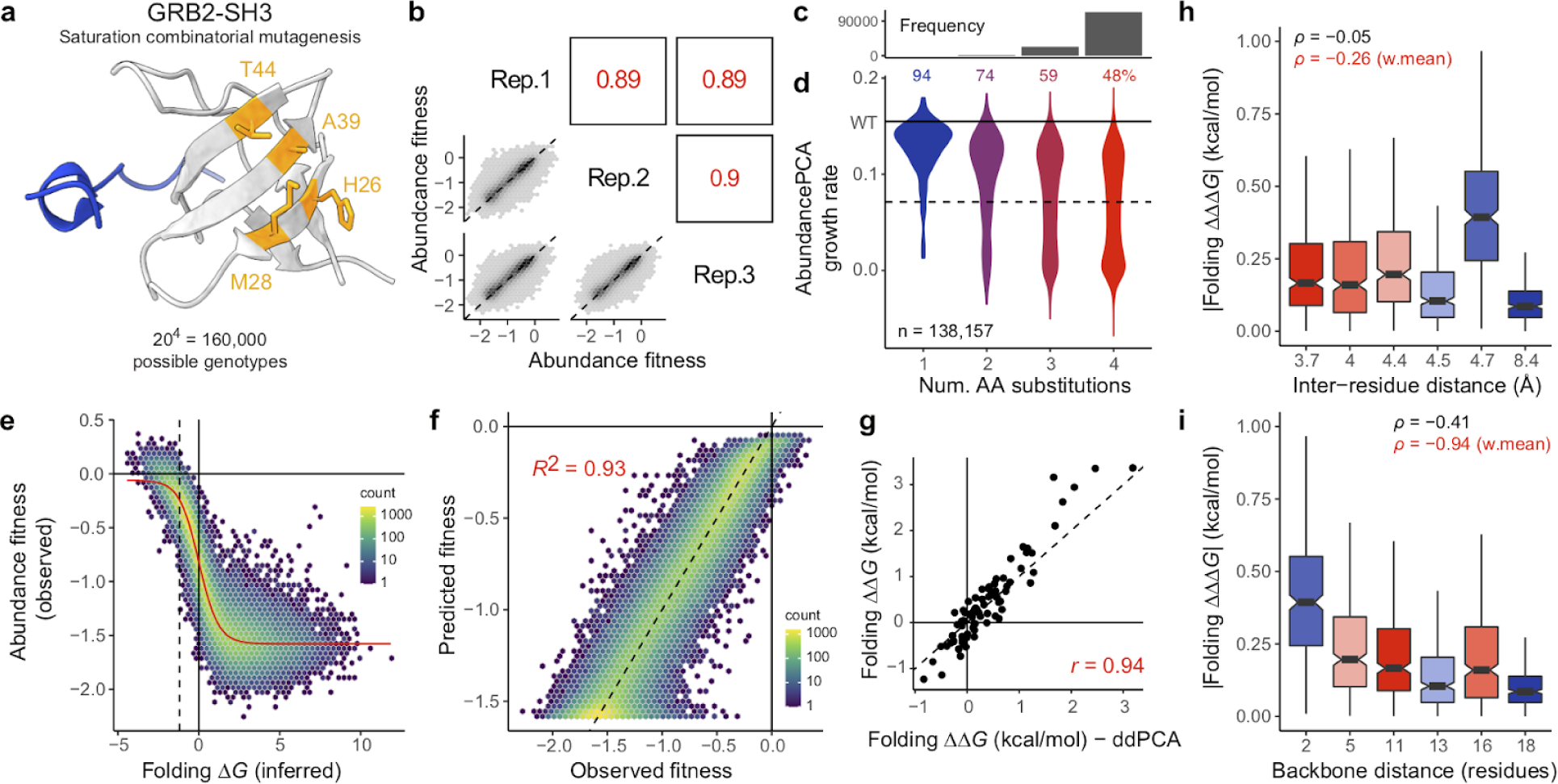
Saturation combinatorial mutagenesis of a protein surface patch. **a**, 3D structure of GRB-SH3 (PDB: 2VWF) indicating four residues targeted for saturation combinatorial mutagenesis (orange) and GAB2 ligand (blue). See also Extended Data Fig. 4. **b**, Scatter plots showing the reproducibility of fitness estimates from AbundancePCA. Pearson’s *r* indicated in red. Rep., replicate. **c**, Histogram showing the number of observed aa variants at increasing Hamming distances from the wild type where the x-axis is shared with panel d. **d**, Violin plot showing distributions of AbundancePCA growth rates inferred from deep sequencing data versus number of amino acid substitutions. The percentage of folded protein variants (predicted fraction folded molecules > 0.5) is shown at each Hamming distance from the wild type. **e**, Non-linear relationship (global epistasis) between observed AbundancePCA fitness and changes in free energy of folding. Thermodynamic model fit shown in red. **f**, Performance of biophysical model including all first- and second-order genetic interaction (energetic coupling) terms/coefficients. **g**, Comparisons of the model-inferred single aa substitution free energy changes to previously reported estimates using GRB2-SH3 ddPCA data ^23^. Pearson’s *r* is shown. **h**, Boxplots showing relationship between folding coupling energy strength and minimal inter-residue side chain heavy atom distance. Boxes are coloured by inter-residue distance. Spearman’s *ρ* is shown for all couplings as well as the weighted mean per residue pair (red). **i**, Relationship between folding coupling energy strength and linear sequence (backbone) distance in number of residues. Boxes are coloured as in panel h. For boxplots in panels h and i: center line, median; box limits, upper and lower quartiles; whiskers, 1.5x interquartile range.

The single mutant effects at these four residues have a larger range than those of the combinatorial library that was designed to conserve fold and function (Fig. 4d). Therefore, when combined in double, triple and quadruple mutants – the most numerous class — the result is a larger fraction of unfolded variants (Fig. 4c-e). A two-state biophysical model including all first- and second-order coefficients provides an excellent fit to the data (*R*^2^ = 0.93, Fig. 4e,f) and inferred folding free energy changes (first-order terms) are highly correlated (Pearson’s *r* = 0.94) with those obtained previously using an independent shallow double mutant library (Fig. 4g, ddPCA). Although the four mutated residues are physically proximal in 3D space, with all except one pair (H26:T44) separated by less than 5 Å (3.8 - 8.4 Å, Fig. 4h), their relative positions in the primary peptide sequence cover a large range (2 - 18 residues, Fig. 4i). There is no relationship between contact distance and folding coupling strength for these residues (Spearman’s *ρ* = -0.05, Fig. 4h), whereas the relationship for backbone distance is significant (Spearman’s *ρ* = -0.41, Fig. 4i). Indeed, backbone distance is almost perfectly correlated with coupling strength when averaging energy terms per residue pair (Spearman’s *ρ* = -0.94, Fig. 4i), with detectable differences even at separation distances greater than 10 residues. The relative position of aa residues in the primary protein sequence is therefore associated with coupling strength independently of their proximity in 3D space.

### Higher order combinatorial mutants are folded and functional

Our experiments identified a large number of GRB2-SH3 genotypes containing higher order combinations of mutations that have high cellular abundance (e.g. 25,564 genotypes containing >5 mutations, Fig. 1f). To further confirm that abundant mutants are correctly folded and functional we performed a third combinatorial mutagenesis experiment in which we also tested the ability of all GRB2-SH3 variants to bind to a peptide ligand using an extensively validated protein-protein interaction assay (BindingPCA ^23,24,32^, Fig. 5a). Recognition of the peptide ligand can only occur if the protein adopts its native conformation ^33^ (Fig. 1b,c). We designed a library consisting of all combinations of 15 single aa substitutions occurring within a 22 aa residue window, avoiding mutations in our original library in binding interface residues (minimal side chain heavy atom distance to the ligand < 5 Å, Extended Data Fig. 4b, see Methods). The library contains 2^15^ (=32,768) variants and shares 6 single aa substitutions with our original 2^34^ library. In total, we obtained binding measurements for 25,967 variants, abundance measurements for 31,936 variants (79% and 97% of the sequence landscape respectively) and the measurements were highly reproducible (Pearson’s *r* > 0.85 and 0.94 for binding and abundance respectively, Fig. 5c-e, Extended Data Fig. 6a-c, see Methods).

**Figure 5.**
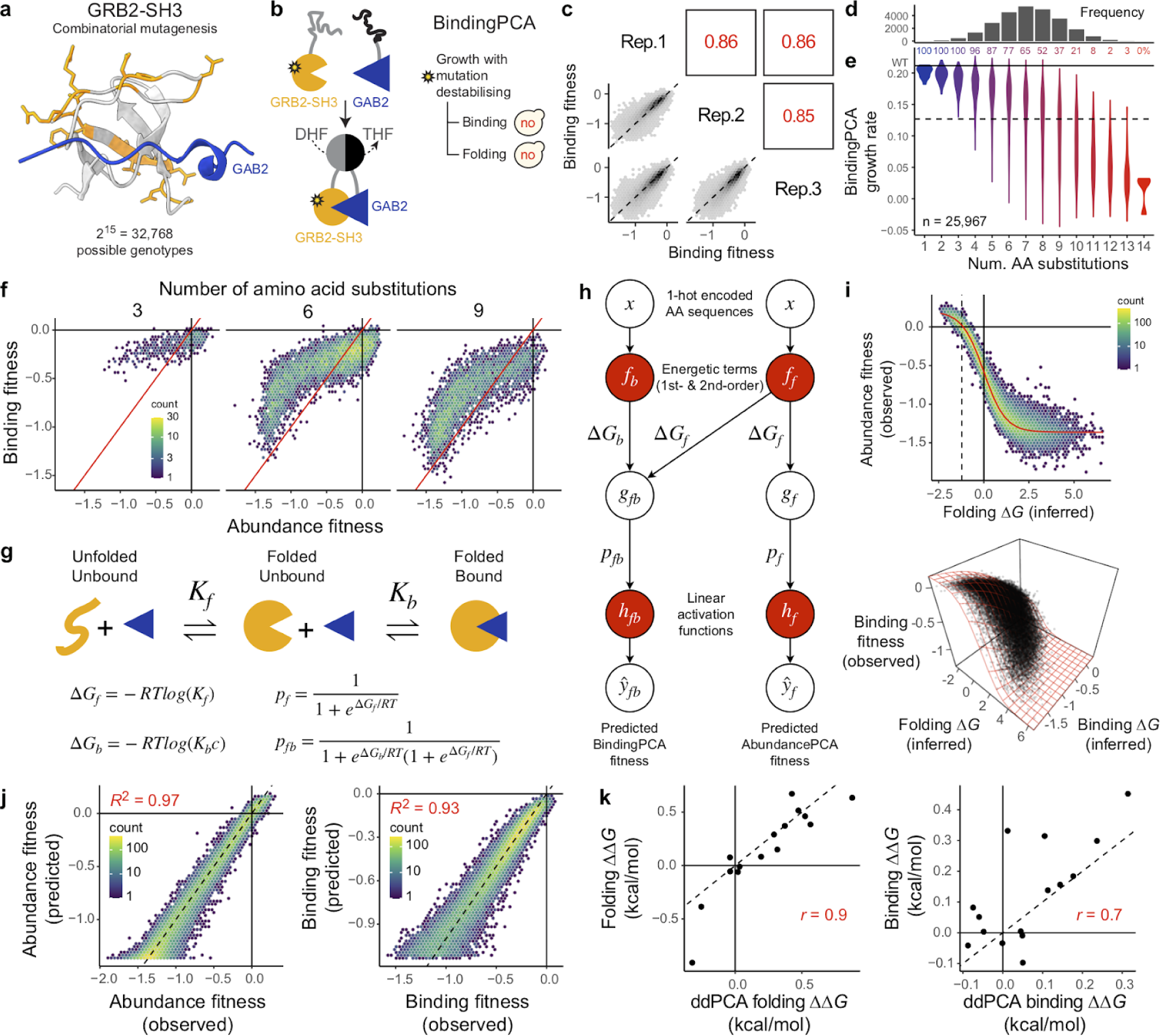
Combinatorial ddPCA reveals that abundant multi-mutants are binding-competent (have a conserved fold). **a,** 3D structure of GRB2-SH3 (PDB: 2VWF) indicating the 15 combinatorially mutated residues in library 3 (orange) and GAB2 ligand (blue). **b**, Overview of BindingPCA on protein of interest (GRB2-SH3) binding to GAB2 ^23^. yes, yeast growth; no, yeast growth defect; DHF, dihydrofolate; THF, tetrahydrofolate. **c**, Scatter plots showing the reproducibility of fitness estimates from BindingPCA. Pearson’s *r* indicated in red. Rep., replicate. **d**, Histogram showing the number of observed aa variants at increasing Hamming distances from the wild type where the x-axis is shared with panel e. **e**, Violin plot showing distributions of BindingPCA growth rates inferred from deep sequencing data versus number of amino acid substitutions. The percentage of bound protein variants (predicted fraction bound molecules > 0.5) is shown at each Hamming distance from the wild type. **f**, 2d density plots comparing abundance and binding fitness of 3rd-(left), 6th-(middle) and 9th-order mutants (right). See Extended Data Fig. 6d for similar plots at all mutant orders. **g**, Three-state equilibrium and corresponding thermodynamic model. Δ*G*_f_, Gibbs free energy of folding; Δ*G*_b_, Gibbs free energy of binding; *K*_f_, folding equilibrium constant; *K*_b_, binding equilibrium constant; *c*, ligand concentration; *p*_f_, fraction folded; *p*_fb_, fraction folded and bound; *g_f_*, non-linear function of Δ*G*_f_; *g_fb_*, non-linear function of Δ*G*_f_ and Δ*G*_b_; R, gas constant; *T*, temperature in Kelvin. **h**, Neural network architecture used to fit thermodynamic model to ddPCA data (bottom, target and output data), thereby inferring the causal changes in free energy of folding and binding associated with single aa substitutions (first-order terms) and pairwise (second-order) interaction terms (top, input values). Δ*G*_f_, Gibbs free energy of folding; *K*_f_, folding equilibrium constant; *p*_f_, fraction folded; *g*, non-linear function of Δ*G*_f_; R, gas constant; *T*, temperature in Kelvin. **i**, Non-linear relationships (global epistasis) between observed AbundancePCA fitness and changes in free energy of folding (top row) or BindingPCA fitness and both free energies of binding and folding (bottom row). Thermodynamic model fit shown in red. **j**, Performance of models fit to ddPCA data. *R*^2^ is the proportion of variance explained. **k**, Comparisons of the model-inferred single aa substitution free energy changes to previously reported estimates using GRB2-SH3 ddPCA data ^23^. Pearson’s *r* is shown.

Plotting the changes in binding against the changes in abundance for 3rd-, 6th- and 9th-order variants reveals that most mutations altering binding also alter the concentration of the isolated domain, consistent with previous results and the expectation that changes in protein stability are a major cause of mutational effects on binding ^23,24,34^ (Fig. 5f, Extended Data Fig. 6d). Most importantly, however, the majority of higher order mutants that have high abundance scores also bind the GAB2 ligand, indicating that they are correctly folded (Fig. 5f, Extended Data Fig. 6d). For example, 4% (204/4,805) of variants containing 9 mutations have abundance indistinguishable from that of the WT protein (nominal *P* > 0.05) and 96% (177/184) of these also bind the ligand (predicted fraction bound molecules > 0.5). The vast majority of abundant higher order GRB2-SH3 mutants are thus correctly folded.

### Accurate multi-phenotype genetic prediction

The exceptionally large number of genetic backgrounds in which both single and double aa mutant effects were measured for these two related molecular phenotypes is a rich source of data for biophysical modeling. First, considering only the abundance phenotype, we observe that an additive two-state thermodynamic model – with unfolded and folded energetic states – outperforms a linear model when evaluated on held out variants (*R*^2^ = 0.93 versus 0.87, Extended Data Fig. 5a,b). In order to attain similar predictive performance as the first-order biophysical model requires including both second- and third-order genetic interaction terms in the linear model (Extended Data Fig. 5c-d), representing a massive increase in model complexity (715 versus 16 i.e. >40-fold more parameters). This apparent greater complexity of inappropriate models that use many specific pairwise and higher order genetic interaction terms to capture global non-linearities in data (global epistasis) has been referred to as ‘phantom epistasis’ ^35^.

Next, extending previous work ^23^, we used a neural network implementation of a three-state equilibrium model ^36^ – with unfolded, folded and bound energetic states (Fig. 5g) – to simultaneously infer the underlying causal free energy changes of both folding and binding (ΔΔ*G*_f_ and ΔΔ*G*_b_), as well as folding and binding energetic couplings (ΔΔΔ*G*_f_ and ΔΔΔ*G*_b_, Fig. 5h). The model fits the data extremely well (Fig. 5i), explaining virtually all of the fitness variance (Fig. 5j), and the inferred folding and binding free energy changes (first-order terms) are well correlated (Pearsons’s *r* = 0.9 and 0.7) with those obtained previously using an independent shallow double mutant library ^23^ (Fig. 5k, ddPCA). This is the first time, to our knowledge, that a large number of folding (ΔΔΔ*G*_f_) and binding (ΔΔΔ*G*_b_) energetic coupling terms have been measured for any protein.

### Allosteric mutations and couplings occur in ligand-proximal residues

We observe that mutational effects on folding energy tend to be larger than those on binding energy (Fig. 6a), recapitulating previous results ^23,24^. Energetic couplings show the same pattern, with folding coupling energies tending to be larger in magnitude than binding energetic couplings (AUC = 0.7, n = 210, *P* = 3.6e-7, two-sided Mann-Whitney U test, Fig. 6b). As none of the mutations in this library occur in the binding interface, any significant effects on binding affinity must be via an allosteric mechanism ^23,24^. Plotting absolute free energy changes against the 3D distance to the ligand reveals a negative correlation as previously reported ^23,24^ (Spearman’s *ρ* = -0.46) with mutations in second-shell residues and residues adjacent (in the primary sequence) to binding interface residues highly enriched for strong allosteric effects on binding affinity (Fig. 6a). Consistent with previous observations ^23,24^, mutations at WT Glycine residues have amongst the strongest effects on binding affinity.

**Figure 6.**
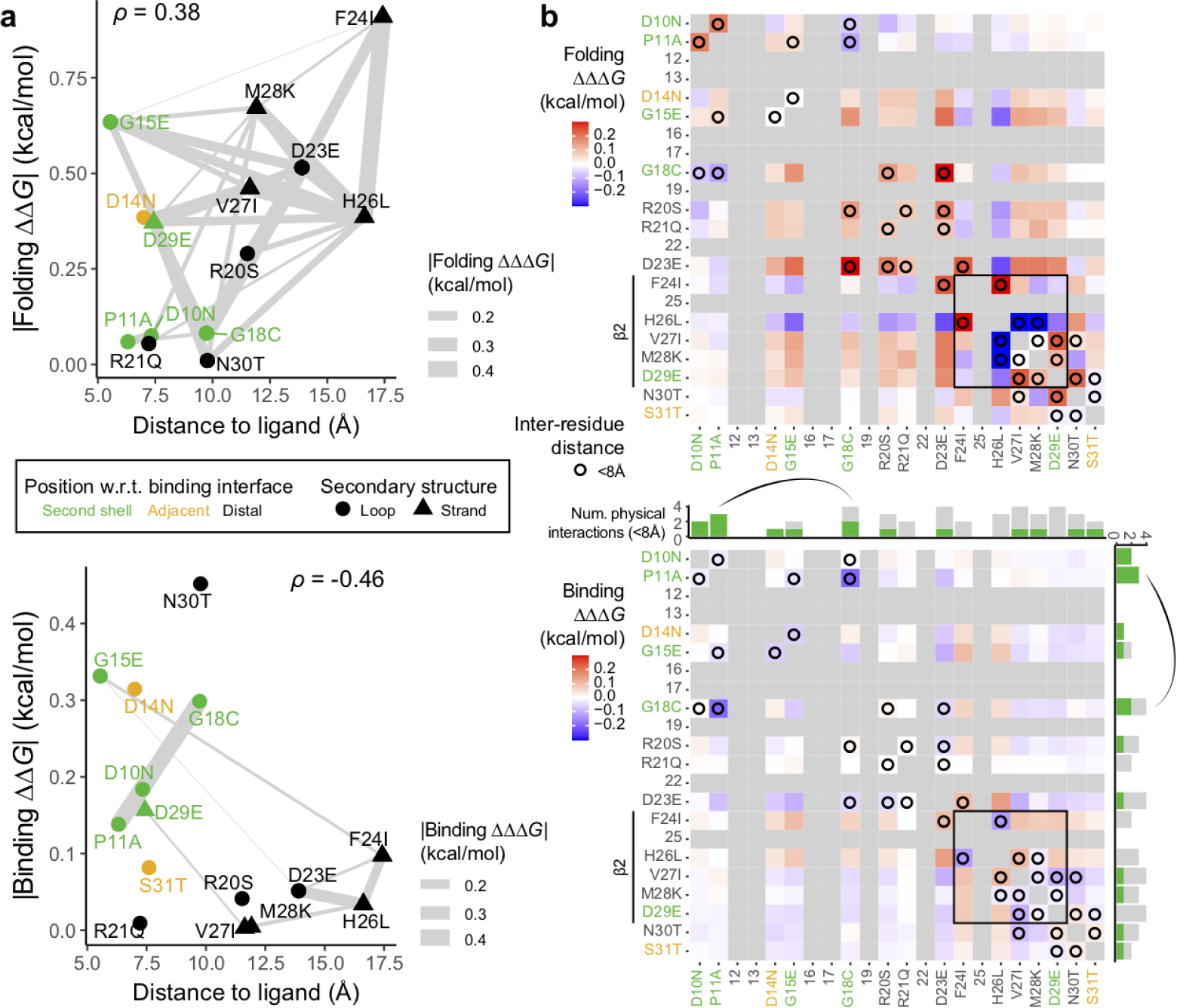
Proximity and connectivity of residues near the binding interface explain energetic effects on binding affinity. **a**, Relationship between the absolute change in free energy of folding (top) and binding (bottom) and minimal side chain heavy atom distance to the ligand. Residues are coloured by their position in the structure relative to the binding interface, triangles indicate beta strand residues and connection lines indicate the strength of energetic couplings between aa pairs (see legend). Spearman’s *ρ* is shown. **b**, Interaction matrix indicating folding (top) and binding coupling terms (bottom) as well as pairwise structural contacts in GRB2-SH3 (PDB: 2VWF, minimal side chain heavy atom distance < 8 Å, black circles). Gray cells indicate missing values (non-mutated residues) and constitutive single aa substitutions are indicated in x- and y-axis labels (see panel a for axis label text color key). Mutation in beta strand residues are indicated and couplings between beta strand residues are boxed. The bar plots above and to the right of the binding interaction matrix indicate the total number of pairwise physical interactions (< 8 Å) involving each residue, with green bars indicating the fraction of interacting partners classified as second shell residues. The strongest binding energetic coupling (P11A:G18C) is indicated.

Finally, whereas the mutations with the strongest folding coupling energies are near-diagonal (closely spaced in the primary sequence), particularly between pairs of residues in the beta strand, the strongest binding coupling in the dataset is an interaction between residues P11 and G18 (Fig. 6b). These two residues are proximal in 3D space (< 8 Å) and constitute one of only two long-range physical contacts between the mutated residues (backbone distance > 5 residues, Fig. 6b), suggesting allosteric energetic couplings are also driven by structural contacts.

## Discussion

By experimentally quantifying protein fold stability in sequence spaces >10^10^ in size we have shown here that the fundamental genetic architecture of at least some proteins is remarkably simple. Biophysical models in which the energetic effects of mutations sum together provide very good prediction of fold stability when tens of mutations are combined. Quantifying the pairwise energetic couplings between mutations further increases predictive power, providing very good performance in high-dimensional genotype spaces. The large number of energetic couplings quantified here reveals important principles about their origins: couplings are strongest between structurally contacting residues and coupling strength also decays along the protein backbone. The genetic architecture of proteins is thus both predictable and understandable.

The biophysical energy models used here are extremely sparse and represent huge data compressions: up to ∼10^8^ (2^34^/34) for the additive models and up to ∼10^7^ (2^34^/596) for the models with energetic couplings. Analyses of previously-published combinatorial protein mutagenesis datasets ^37,38^, mutagenesis of a protein interaction interface ^38^, combinatorial mutagenesis of a tRNA ^39^ and mutagenesis of an alternatively spliced exon ^35^ suggest that this simplicity of genotype-phenotype landscapes is widely observed and likely to be a general principle of macromolecules and their molecular reactions.

Energy models are grounded in our understanding of protein physics and their simplicity and interpretability contrasts with the extreme complexity and lack of mechanistic insight provided by deep neural networks. Predictive energy models are likely to have many applications, including for clinical variant effect interpretation ^40^, pathogen and pandemic forecasting ^41^, and protein engineering for biotechnology ^42,43^. An important challenge moving forward is how to efficiently quantify the free energy changes and energetic couplings for all mutations in proteins of interest. Quantifying mutational effects across diverse genetic backgrounds and homologous sequences may be an efficient way to achieve this.

Our data do not rule out the importance of higher order genetic interactions for protein stability. Rather they show that when global nonlinearities due to cooperative protein folding are properly accounted for and measurements are averaged across genetic backgrounds, first-order and pairwise energetic couplings provide sufficient information for many prediction tasks. An important question to address in future work will be the extent to which higher-order energetic interactions become important in even larger sequence spaces, including in the twilight zone of structurally homologous proteins with very low sequence identity. Quite simple experimental designs should be able to definitely address this question for a diversity of protein folds.

## Methods

### Combinatorial mutagenesis library designs

#### Combinatorial library 1

Library 1 was designed using a computationally efficient greedy strategy to search for the largest number of single aa substitutions which, when combined, preserve both fold and function even in the highest order mutants (Fig. 1b). The algorithm used previously-published ddPCA data and biophysical modeling results for GRB2-SH3, including inferred single aa substitution free energy changes of folding and binding for this protein ^23^. We showed previously that this model – which assumes individual inferred folding and binding free energy changes (ΔΔ*G*_f_ and ΔΔ*G*_b_) combine additively in multi-mutants – accurately predicts the effects of double aa substitutions ^23^. This same additive model was therefore used to make predictions about the energetic and phenotypic effects of higher order mutants explored in the greedy search.

First, the set of candidate single aa mutations was restricted to those with confident free energy changes defined as those with 95% confidence intervals < 1 kcal/mol and whose effects were measured in at least 20 genetic backgrounds (i.e. double aa mutations). Candidate mutations were further restricted to those reachable by single nucleotide substitutions in the wild-type sequence in order to simplify synthesis of the resulting combinatorial mutagenesis library. The algorithm begins from an arbitrary starting mutation and iteratively selects additional mutations at other residue positions until all residues in the protein have been mutated. The heuristic works by selecting additional mutations at each step that maximize the fold and function of the current highest order mutant combination i.e. geometric mean of model predicted AbundancePCA and BindingPCA growth rates. This procedure is then repeated for all possible starting mutations.

In order to visualize and compare the resulting solutions we also simulated the median AbundancePCA and BindingPCA growth rates of all candidate combinatorial libraries, calculated using a random sample of 10,000 variants. Although the algorithm is not guaranteed to produce the optimal solution at each Hamming distance from the wild-type sequence, the greedy approach nevertheless achieves solutions in which both phenotypes are predicted to be preserved in variants with greater than 30 mutations (Extended Data Fig. 1b), beyond which one or both phenotypes are lost. Defining viable libraries as those preserving both molecular phenotypes above 70% of the maximal value (i.e. geometric mean of simulated median AbundancePCA and BindingPCA growth rates) resulted in the largest candidate combinatorial library consisting of all combinations of 34 single aa mutations (Fig. 1, Extended Data Fig. 1b-d).

#### Combinatorial library 2

We clustered the contact map (minimal side chain heavy atom distance < 5 Å) comprising all GRB2-SH3 surface residues (RSASA ≥ 0.25) existing in secondary structure elements (Extended Data Fig. 4) and selected the following four physically proximal residues for saturation combinatorial mutagenesis: H26, M28, A39, T44 (see Fig. 4).

#### Combinatorial library 3

This library was designed to include all combinations of 15 single aa substitutions with mild effects (within one third of the AbundancePCA fitness interquartile range of the wild type ^23^) in close proximity in the primary sequence and reachable by single nucleotide substitutions while avoiding mutations in binding interface residues (minimal side chain heavy atom distance to the ligand < 5 Å). We used a sliding window approach to determine the number of candidate mutant residues in stretches of 20, 21 and 22 consecutive residues in GRB2-SH3 (Extended Data Fig. 4b). Only one window with a width of 22 aa (starting at residue position 10) includes 15 candidate positions (Extended Data Fig. 4b). The final library consisted of all combinations of the following randomly selected candidate mutations at these positions: D10N, P11A, D14N, G15E, G18C, R20S, R21Q, D23E, F24I, H26L, V27I, M28K, D29E, N30T, S31T (see Fig. 5).

### Mutagenesis library construction and selection assays

#### Media and buffers used

- LB: 10 g/L Bacto-tryptone, 5 g/L Yeast extract, 10 g/L NaCl. Autoclaved 20 min at 120°C.
- YPDA: 20 g/L glucose, 20 g/L Peptone, 10 g/L Yeast extract, 40 mg/L adenine sulphate. Autoclaved 20 min at 120°C.
- SORB: 1 M sorbitol, 100 mM LiOAc, 10 mM Tris pH 8.0, 1 mM EDTA. Filter sterilized (0.2 mm Nylon membrane, ThermoScientific).
- Plate mixture: 40% PEG3350, 100 mM LiOAc, 10 mM Tris-HCl pH 8.0, 1 mM EDTA pH 8.0. Filter sterilised.
- Recovery medium: YPD (20 g/L glucose, 20 g/L Peptone, 10 g/L Yeast extract) + 0.5 M sorbitol. Filter sterilised.
- SC -URA: 6.7 g/L Yeast Nitrogen base without amino acid, 20 g/L glucose, 0.77 g/L complete supplement mixture drop-out without uracil. Filter sterilised.
- SC -URA/MET/ADE: 6.7 g/L Yeast Nitrogen base without amino acid, 20 g/L glucose, 0.74 g/L complete supplement mixture drop-out without uracil, adenine and methionine. Filter sterilised.
- Competition medium: SC –URA/MET/ADE + 200 ug/mL methotrexate (MERCK LIFE SCIENCE), 2% DMSO.
- DNA extraction buffer: 2% Triton-X, 1% SDS, 100mM NaCl, 10mM Tris-HCl pH8, 1mM EDTA pH8.

#### Plasmid construction

GRB2 mutagenesis plasmid pGJJ286: wt GRB2-SH3 was digested from pGJJ046 (described in ^23^) with the restriction enzymes AvrII and HindIII and cloned into the digested plasmid pGJJ191 (described in ^24^) using T4 ligase (New England Biolabs). AbundancePCA (aPCA) pGJJ046 and pGJJ045 plasmids and BindingPCA (bPCA) pGJJ034 and pGJJ001 plasmids were previously described in ^23^.

#### Libraries construction

Libraries were constructed in two steps. First, an IDT primer containing the chosen combination of mutations was assembled by Gibson into the mutagenesis plasmid pGJJ286. Libraries were then cloned into the yeast plasmids aPCA (pGJJ045) and bPCA (pGJJ001) by digestion/ligation. For the first step the libraries into the mutagenesis plasmid were assembled by Gibson reaction (in house preparation) of two fragments. The vector fragment was obtained by PCR amplification of pGJJ286 with the oligos shown in the Supplementary Tables 1 and 2, incubated with DpnI to remove the template and gel purified using QIAquick gel extraction kit (Qiagen). The insert fragment was obtained by mixing equimolar amounts of IDT mutation primer (Supplementary Tables 1 and 2) and a reverse elongation primer (Supplementary Tables 1 and 2) and incubating for 1 cycle of annealing/extension with Q5 polymerase (New England Biolabs). dsDNA product was then incubated with ExoSAP-IT (Applied Biosystem) to get rid of the remaining ssDNA and purified with MinElute columns (Qiagen). 100ng of vector in a molar ratio 1:5 with the insert were incubated for 3h at 50°C with a gibson mix 2x prepared in house. The reaction was desalted by dialysis with membrane filters (MF-Millipore) for 1h and concentrated 4X using SpeedVac concentrator (Thermo Scientific). DNA was then transformed into NEB 10β High-efficiency Electrocompetent E.Coli. Cells were allowed to recover in SOC medium (NEB 10β Stable Outgrowth Medium) for 30 minutes and later transferred to LB medium with spectinomycin overnight. A fraction of cells was also plated into spectinomycin+LB+agar plates to estimate the total number of transformants. 100 mL of each saturated *E. coli* culture were harvested next morning to extract the mutagenesis plasmid library using the QIAfilter Plasmid Midi Kit (QIAGEN). To obtain the final libraries into the yeast plasmids, libraries in pGJJ286 plasmid were digested with NheI and HindIII, gel purified (MinElute Gel Extraction Kit, QIAGEN) and cloned into pGJJ045 or pGJJ034 digested plasmids with T4 ligase (New England Biolabs) by temperature-cycle ligation following manufacturer instructions, 67 fmol of backbone and 200 fmol of insert in a 33.3 uL reaction. The ligation was desalted by dialysis using membrane filters for 1h, concentrated 4X using a SpeedVac concentrator (Thermo Scientific) and transformed into NEB 10β High-efficiency Electrocompetent E. coli cells.

#### Methotrexate yeast selection assay

The yeast selection assay was described in ^23^. The high-efficiency yeast transformation protocol described below (adjusted to a pre-culture of 200 mL of YPDA) was scaled up or down depending on the number of transformants for each library (Supplementary Table 2). Three independent pre-cultures of BY4742 were grown in 20 mL standard YPDA at 30°C overnight. The next morning, the cultures were diluted into 200 mL of pre-wormed YPDA at an OD_600nm_ =

0.3 and incubated at 30°C for 4 hours. Cells were then harvested and centrifuged for 5 minutes at 3,000g, washed with sterile water and SORB medium, resuspended in 8.6 mL of SORB and incubated at room temperature for 30 minutes. After incubation, 175 μL of 10mg/mL boiled salmon sperm DNA (Agilent Genomics) and 3.5 μg of plasmid library were added to each tube of cells and mixed gently. 35 mL of Plate Mixture were added to each tube to be incubated at room temperature for an additional 30 minutes. 3.5 mL of DMSO was added to each tube and the cells were then heat shocked at 42°C for 20 minutes (inverting tubes from time to time to ensure homogenous heat transfer). After heat shock, cells were centrifuged and re-suspended in ∼50 mL of recovery media and allowed to recover for 1 hour at 30°C. Cells were then centrifuged, washed with SC-URA medium and re-suspended in 200mL SC -URA. 10 uL were plated on SC -URA Petri dishes and incubated for ∼48 hours at 30°C to measure the transformation efficiency. The independent liquid cultures were grown at 30°C for ∼48 hours until saturation. Saturated cells were diluted again to OD_600nm_ = 0.1 in SC -URA/MET/ADE media and allowed to grow 4 generations until OD_600nm_ = 1.6 at 30°C and 200rpm. A fraction of the culture was then used to inoculate 200mL of competition media containing methotrexate at a starting OD_600nm_ = 0.05, and the rest was harvested and pellets frozen and stored as INPUT. Cells in competition media were allowed to grow for 3-5 generations (Supplementary Table 2), collected and frozen and stored as OUTPUT.

#### DNA extractions and plasmid quantification

The DNA extraction protocol used was described in ^23^. Protocol below is for 100 mL of harvested culture at OD_600nm_ ∼ 1.6. Protocols were scaled up or down depending on the library (Supplementary Table 2). Cell pellets (one for each experiment input/output replicate) were re-suspended in 1 mL of DNA extraction buffer, frozen by dry ice-ethanol bath and incubated at 62°C water bath twice. Subsequently, 1 mL of Phenol/Chloro/Isoamyl 25:24:1 (equilibrated in 10mM Tris-HCl, 1mM EDTA, pH8) was added, together with 1 g of acid-washed glass beads (Sigma Aldrich) and the samples were vortexed for 10 minutes. Samples were centrifuged at RT for 30 minutes at 4,000 rpm and the aqueous phase was transferred into new tubes. The same step was repeated twice. 0.1 mL of NaOAc 3M and 2.2 mL of pre-chilled absolute ethanol were added to the aqueous phase. The samples were gently mixed and incubated at -20°C at least for 30 minutes. After that, they were centrifuged for 30 min at full speed at 4°C to precipitate the DNA. The ethanol was removed and the DNA pellet was allowed to dry overnight at RT. DNA pellets were resuspended in 0.6 mL TE 1X and treated with 5 uL of RNaseA (10mg/mL, Thermo Scientific) for 30 minutes at 37°C. To desalt and concentrate the DNA solutions, QIAEX II Gel Extraction Kit was used (50 µL of QIAEX II beads, QIAGEN). The samples were washed twice with PE buffer and eluted twice by 125 µL of 10 mM Tris-HCI buffer, pH 8.5. Finally, plasmid concentrations in the total DNA extract (that also contained yeast genomic DNA) were quantified by qPCR using the primer pair oGJJ152-oGJJ153, that binds to the ori region of the plasmids.

#### Sequencing library preparation

This was shown in ^23^. Briefly, the sequencing libraries were constructed in two consecutive PCR reactions. The first PCR (PCR1) was designed to amplify the mutated protein of interest and to increase the nucleotide complexity of the first sequenced bases by introducing frame-shift bases between the adapters and the sequencing region of interest (Supplementary Tables 1 and 2). The second PCR (PCR2) was necessary to add the remainder of the Illumina adapter and demultiplexing indexes. PCR2 reactions were run for each sample independently using Hot Start High-Fidelity DNA Polymerase. In this second PCR the remaining parts of the Illumina adapters were added to the library amplicon. The forward primer (5’ P5 Illumina adapter) was the same for all samples (GJJ_1J), while the reverse primer (3’ P7 Illumina adapter) differed by the barcode index (Supplementary Table 3) to be subsequently pooled together and demultiplex them after deep sequencing. All samples were pooled in an equimolar ratio and gel purified using the QIAEX II Gel Extraction Kit. The purified amplicon library pools were subjected to Illumina 150bp paired-end Nextseq500 sequencing at the CRG Genomics Core Facility.

### Sequencing data processing

FastQ files from paired-end sequencing of all AbundancePCA and BindingPCA experiments were processed with DiMSum v1.3 ^44^ using default settings with minor adjustments: https://github.com/lehner-lab/DiMSum. Supplementary Table 4 contains DiMSum fitness estimates and associated errors for all experiments. Experimental design files and command-line options required for running DiMSum on these datasets are available on GitHub (https://github.com/lehner-lab/archstabms). Variants with less than 10 Input read counts in any replicate were discarded (“fitnessMinInputCountAll” option) i.e. only variants observed in all replicates above this threshold were retained. For library 1 we also included fitness estimates that derived from a subset of replicates whose Input read counts exceeded this threshold (“fitnessMinInputCountAny” option, see Fig. 1).

In view of the length and complexity of mutagenesis library 1 we also included a wild-type-only sample for sequencing using pGJJ046 as template in order to derive empirical estimates of sequencing error rates. The FastQ file for this sample was processed identically to those of the replicate Input/Output samples in the first pass analysis with DiMSum with permissive base quality thresholds (“vsearchMinQual = 5” and “vsearchMaxee = 1000”). Read counts for all variants were then adjusted by subtracting the expected number of sequencing errors derived from the wild-type-only sample and proportional to the total sequencing library size of each sample. Finally, fitness estimates and associated errors for library 1 were then obtained from the resulting corrected variant counts with DiMSum (“countPath” option).

### Thermodynamic modeling with MoCHI

We used MoCHI (https://github.com/lehner-lab/MoCHI) to fit all thermodynamic models to combinatorial DMS data using default settings with minor adjustments. The software is based on our previously described genotype-phenotype modeling approach ^23^ with additional functionality and improvements for ease-of-use and flexibility ^24^. Models fit to shallow (ddPCA) libraries and used in the analyses described in this work (e.g. combinatorial mutagenesis library designs) were obtained using the original software implementation ^23^.

We model protein folding as an equilibrium between two states: unfolded (u) and folded (f), and protein binding as an equilibrium between three states: unfolded and unbound (uu), folded and unbound (fu), and folded and bound (fb). We assume that the probability of the unfolded and bound state (ub) is negligible and free energy changes of folding and binding are additive i.e. the total binding and folding free energy changes of an arbitrary variant relative to the wild-type sequence is simply the sum over residue-specific energies corresponding to all constituent single amino acid substitutions.

We configured MoCHI parameters to specify a neural network architecture consisting of additive trait layers (free energies) for each biophysical trait to be inferred (folding or folding and binding for AbundancePCA or BindingPCA, respectively), as well as one linear transformation layer per observed phenotype. The specified non-linear transformations “TwoStateFractionFolded” and “ThreeStateFractionBound” derived from the Boltzmann distribution function relate energies to proportions of folded and bound molecules respectively (see Fig. 2a and Fig. 5g,h). The target (output) data to fit the neural network comprises fitness scores for the wild type and aa substitution variants of all mutation orders. The inclusion of both first- and second-order (pairwise energetic coupling) model coefficients in the models was specified using the “max_interaction_order” option.

A random 30% of aa substitution variants of all mutation orders was held out during model training, with 20% representing the validation data and 10% representing the test data. Validation data was used to evaluate training progress and optimize hyperparameters (batch size). Optimal hyperparameters were defined as those resulting in the smallest validation loss after 100 training epochs. Test data was used to assess final model performance.

MoCHI optimizes the parameters θ of the neural network using stochastic gradient descent on a loss function ℒ[θ] based on a weighted and regularized form of mean absolute error:

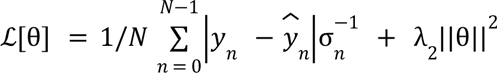

where *y_n_* and σ*_n_* are the observed fitness score and associated standard error respectively for variant *n*, *y*^^^*_n_* is the predicted fitness score, *N* is the batch size and λ_2_ is the *L*_2_ regularization penalty. In order to penalize very large free energy changes (typically associated with extreme fitness scores) we set λ_2_ to 10^-6^ representing light regularization. The mean absolute error is weighted by the inverse of the fitness error (σ^-1^*_n_*) in order to downweight the contribution of less confidently estimated fitness scores to the loss. Furthermore, in order to capture the uncertainty in fitness estimates, the training data was replaced with a random sample from the fitness error distribution of each variant. The validation and test data was left unaltered.

Models were trained with default settings i.e. for a maximum of 1000 epochs using the Adam optimization algorithm with an initial learning rate of 0.05 (except for library 1 for which we used an initial learning rate of 0.005). MoCHI reduces the learning rate exponentially (γ = 0. 98) if the validation loss has not improved in the most recent ten epochs compared to the preceding ten epochs. In addition, MoCHI stops model training early if the wild-type free energy terms over the most recent ten epochs have stabilized (standard deviation ≤10^−3^).

Free energies are calculated directly from model parameters as follows: Δ*G_b_* = θ*_b_RT* and Δ*G_f_* = θ *_f_RT*, where *T* = 303 K and *R* = 0.001987 kcalK^-1^mol^-1^. We estimated the confidence intervals of model-inferred free energies using a Monte Carlo simulation approach. The variability of inferred free energy changes was calculated between ten separate models fit using data from [1] independent random training-validation-test splits and [2] independent random samples of fitness estimates from their underlying error distributions. Confident inferred free energy changes are defined as those with Monte Carlo simulation derived 95% confidence intervals < 1 kcal/mol. Supplementary Table 5 contains inferred binding and folding free energy changes and energetic couplings from all second-order models.

### Linear model to predict energetic coupling strength

We built a linear model to predict energetic coupling strength (absolute value of energetic coupling terms) from twelve features (see Fig. 3e), 5 distance metrics for residue pairs or positions thereof in the protein structure: backbone distance (linear 1D distance separating residue pairs along the primary aa sequence), inter-residue distance (minimal side-chain heavy atom distance in 3D space), number of core residues (0, 1, or both residues in the pair with RSASA < 0.25), number of binding interface residues (0, 1, or both with minimal side chain heavy atom distance to the ligand < 5 Å), number of beta sheet residues (0, 1, or both in beta strands) and 7 features describing the number of chemical bonds or interactions between the atoms of pairs of residues as calculated using the GetContacts software tool (https://getcontacts.github.io/): backbone to backbone hydrogen bonds, side-chain to backbone hydrogen bonds, side-chain to side-chain hydrogen bonds, Pi-cation interactions, Pi-stacking interactions, Salt bridge interactions and van der Waals interactions. Before running GetContacts we used PyMOL to fill missing hydrogens (“h_add” command), FoldX ^45^ to restore the wild-type Proline at position 54 that is mutated in the reference crystal structure (PDB: 2VWF, “PositionScan” command) and removed GAB2 ligand atoms. The training dataset comprised energetic couplings inferred from library 1 and the test set comprised independently inferred energetic couplings from library 3 (see Fig. 3f).

## Acknowledgements

This work was funded by European Research Council (ERC) Advanced grant (883742), the Spanish Ministry of Science and Innovation (LCF/PR/HR21/52410004, EMBL Partnership, Severo Ochoa Centre of Excellence), the Bettencourt Schueller Foundation, the AXA Research Fund, Agencia de Gestio d’Ajuts Universitaris i de Recerca (AGAUR, 2017 SGR 1322), and the CERCA Program/Generalitat de Catalunya. A.J.F. was funded by a Ramón y Cajal fellowship (RYC2021-033375-I) financed by the Spanish Ministry of Science and Innovation (MCIN/AEI/10.13039/501100011033) and the European Union (NextGenerationEU/PRTR). We thank all members of the Lehner Lab for helpful discussions and suggestions.

## Author contributions

B.L. and A.J.F. conceived the project motivated in part by preliminary unpublished analyses of evolutionary couplings performed by J.M.S. B.L., A.J.F. and A.M.-A. designed the combinatorial mutagenesis libraries and experiments. A.M. and C.H.-C. performed the experiments with help from A.J.F. A.J.F. led the data analysis with help from A.M.-A. A.J.F. and B.L. wrote the manuscript with input from A.M.-A. and C.H.-C.

## Additional information

Supplementary Information is available for this paper. Correspondence and requests for materials should be addressed to A.J.F and B.L.

## Data availability

All DNA sequencing data have been deposited in the Gene Expression Omnibus with accession number GSE246322: https://www.ncbi.nlm.nih.gov/geo/query/acc.cgi?acc=GSE246322. All fitness measurements and free energies are provided in Supplementary tables 4 and 5.

## Code availability

Source code for fitting thermodynamic models (MoCHI) is available at https://github.com/lehner-lab/MoCHI. Source code for all downstream analyses, including DiMSum and MoCHI configuration files and to reproduce all figures described here is available at https://github.com/lehner-lab/archstabms.

## Extended Data Figures

**Extended Data Figure 1.**
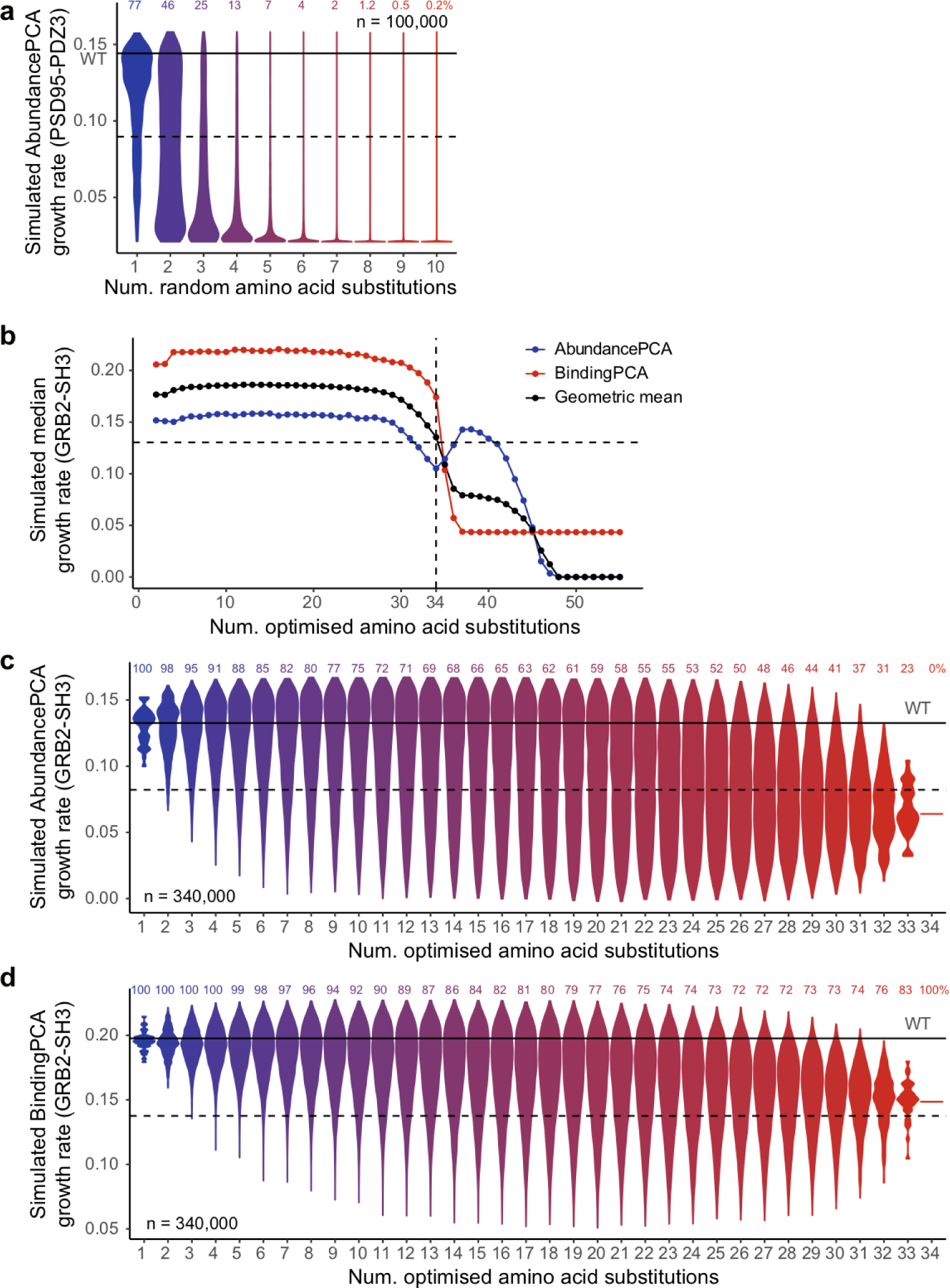
Combinatorial mutagenesis library 1 design and simulations. **a**, Violin plot showing distributions of simulated AbundancePCA growth rates (assuming additivity of individual inferred folding free energy changes ^23^) versus number of random amino acid substitutions (n = 100,000) for PSD95-PDZ3. Violins are scaled to have the same maximum width. **b**, Simulated median AbundancePCA/BindingPCA growth rates of optimal combinatorial libraries of increasing maximum amino acid Hamming distances from the wild type. The horizontal dashed line indicates the 70th percentile of the maximal geometric mean (black). The vertical dashed line indicates the number of amino acid substitutions selected (n = 34) for the synthesized combinatorial mutagenesis library 1. **c**, Violin plot showing simulated distributions of AbundancePCA growth rates versus number of amino acid substitutions for combinatorial mutagenesis library 1. **d**, similar to panel c, but showing simulated distributions for BindingPCA growth rates. In panels c and d, the percentage of folded and bound protein variants (predicted fraction folded or bound molecules > 0.5) is shown at each Hamming distance from the wild type.

**Extended Data Figure 2.**
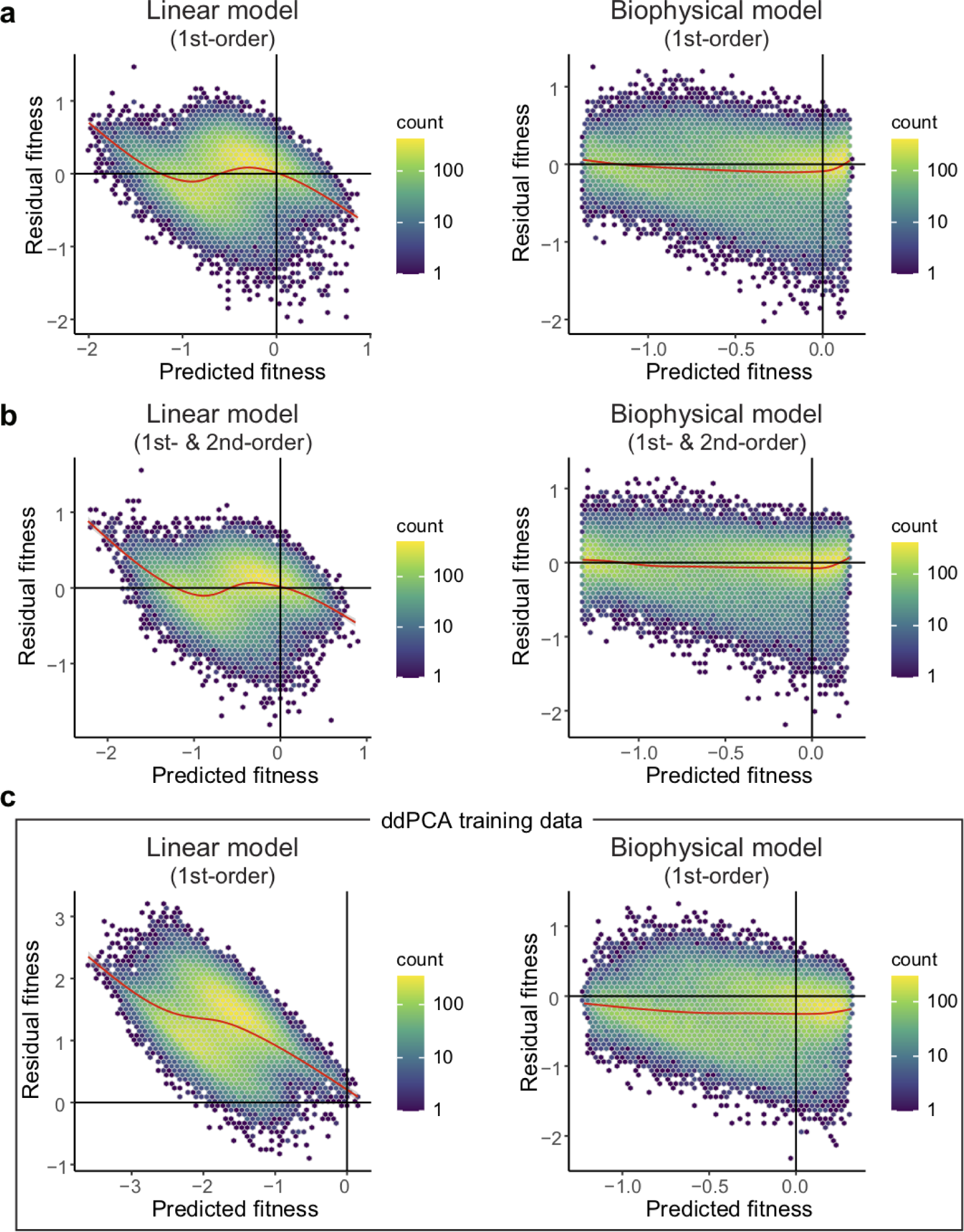
Residuals versus fitted values for linear and biophysical models fit to AbundancePCA data from combinatorial library 1. **a**, Residual fitness (observed - predicted) versus predicted fitness for first-order linear models (left) and first-order biophysical models (right) evaluated on GRB2-SH3 combinatorial AbundancePCA data (combinatorial library 1, see Fig. 1). The smoothed conditional mean (generalized additive model) is shown in red. **b**, Similar to panel a except models include all first- and second-order genetic interaction (energetic coupling) terms/coefficients. **c**, Similar to panel a except results are shown for models that were trained on GRB2-SH3 ddPCA data consisting of single and double aa substitutions only ^23^.

**Extended Data Figure 3.**
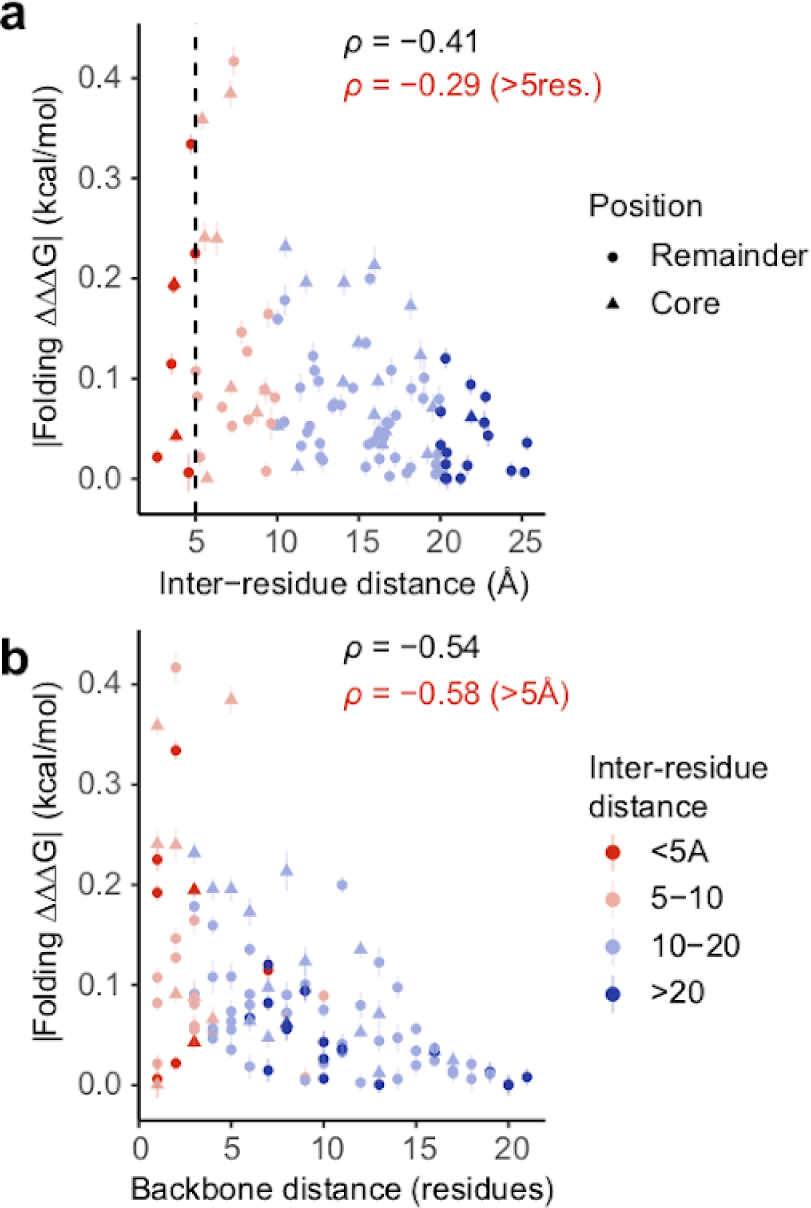
Structural determinants of energetic couplings inferred from ddPCA data from combinatorial library 3. **a**, Relationship between folding coupling energy strength and minimal inter-residue side chain heavy atom distance for combinatorial library 3 (see Fig. 5). Error bars indicate 95% confidence intervals from a Monte Carlo simulation approach (*n* = 10 experiments). Points are coloured by binned inter-residue distances (see legend in panel b). Spearman’s *ρ* is shown for all couplings as well as those involving pairs of residues separated by more than five residues in the primary sequence (red). Core residues are indicated as triangles. **b**, Relationship between folding coupling energy strength and linear sequence (backbone) distance in number of residues.

**Extended Data Figure 4.**
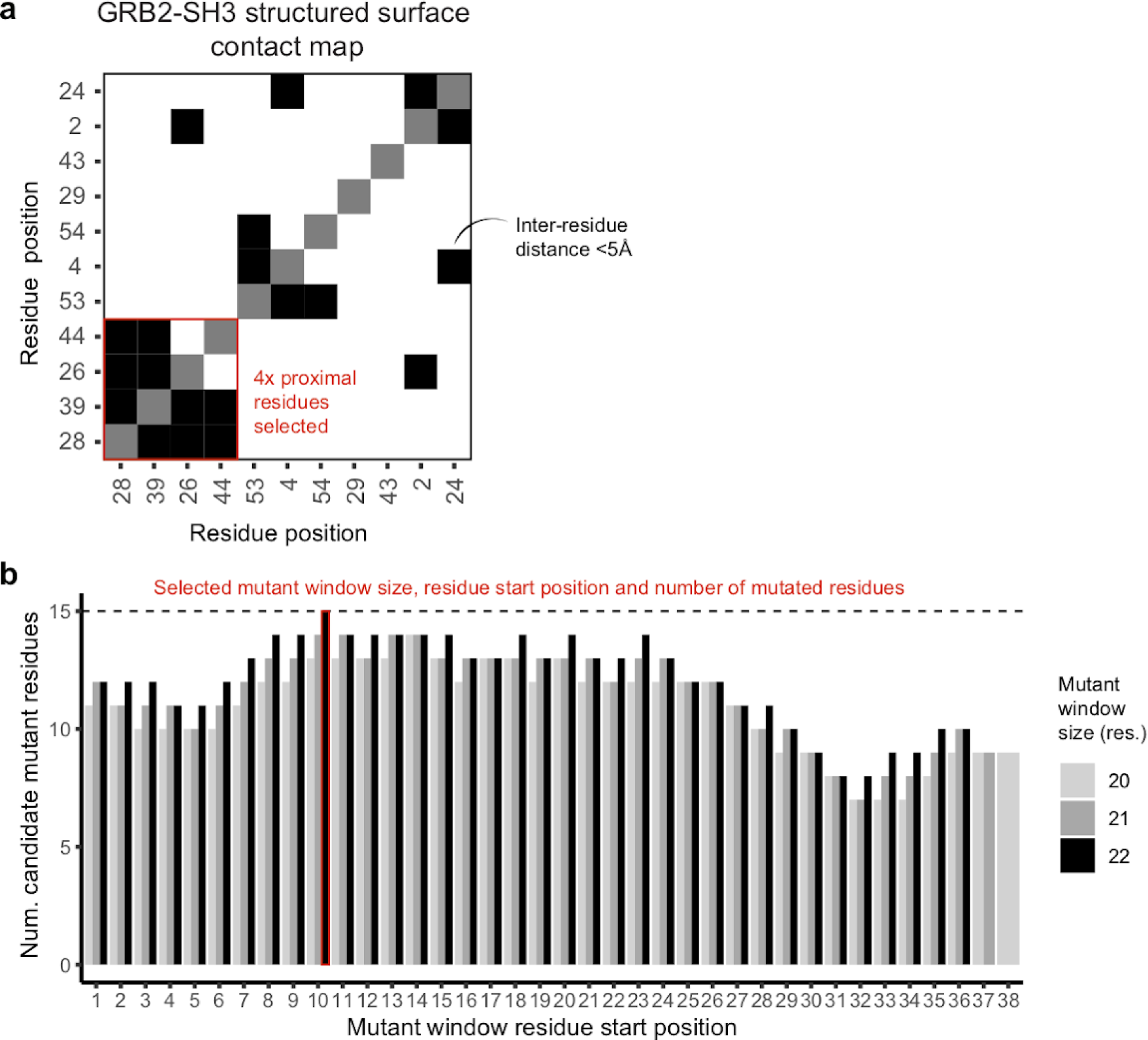
Design of combinatorial mutagenesis libraries 2 and 3. **a**, Clustered heat map showing structural contacts (minimal side chain heavy atom distance < 5 Å) between all GRB2-SH3 surface residues (RSASA ≥ 0.25) existing in secondary structure elements. The four highlighted residues are all physically proximal and were selected as the targets for library 2 saturation combinatorial mutagenesis (see Fig. 4). **b**, Bar plot indicating the number of candidate mutant residues in stretches of 20, 21 and 22 consecutive residues in GRB2-SH3 used to design mutagenesis library 3. Candidate mutations were defined as single aa substitutions with mild effects (within one third of the AbundancePCA fitness interquartile range of the wild type ^23^) in close proximity in the primary sequence and reachable by single nucleotide substitutions while avoiding mutations in binding interface residues (minimal side chain heavy atom distance to the ligand < 5 Å). The selected mutant window size (22 aa residues), residue start position (10) and number of mutated residues (15) is indicated. The final library consisted of all combinations of the following randomly selected candidate mutations at these 15 positions: D10N, P11A, D14N, G15E, G18C, R20S, R21Q, D23E, F24I, H26L, V27I, M28K, D29E, N30T, S31T (see Fig. 5).

**Extended Data Figure 5.**
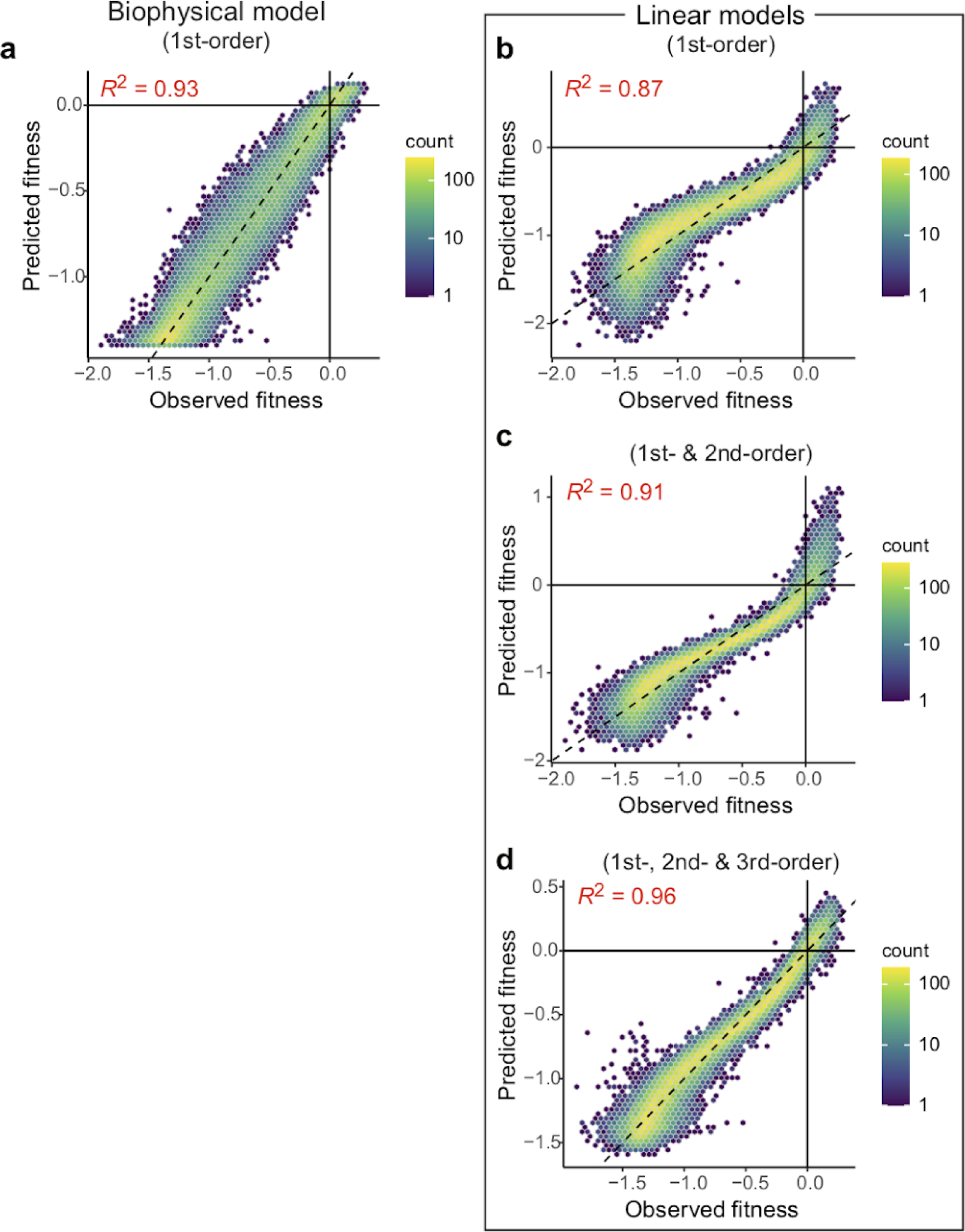
Performance of models fit to AbundancePCA data from combinatorial library 3. **a**, Performance of first-order 2-state biophysical model (folded and unfolded states) fit to AbundancePCA data from combinatorial library 3 (see Fig. 5). **b-d**, Performance of first- (panel b), second- (panel c) and third-order (panel d) linear models fit to AbundancePCA data from combinatorial library 3 (see Fig. 5). *R*^2^ is the proportion of variance explained.

**Extended Data Figure 6.**
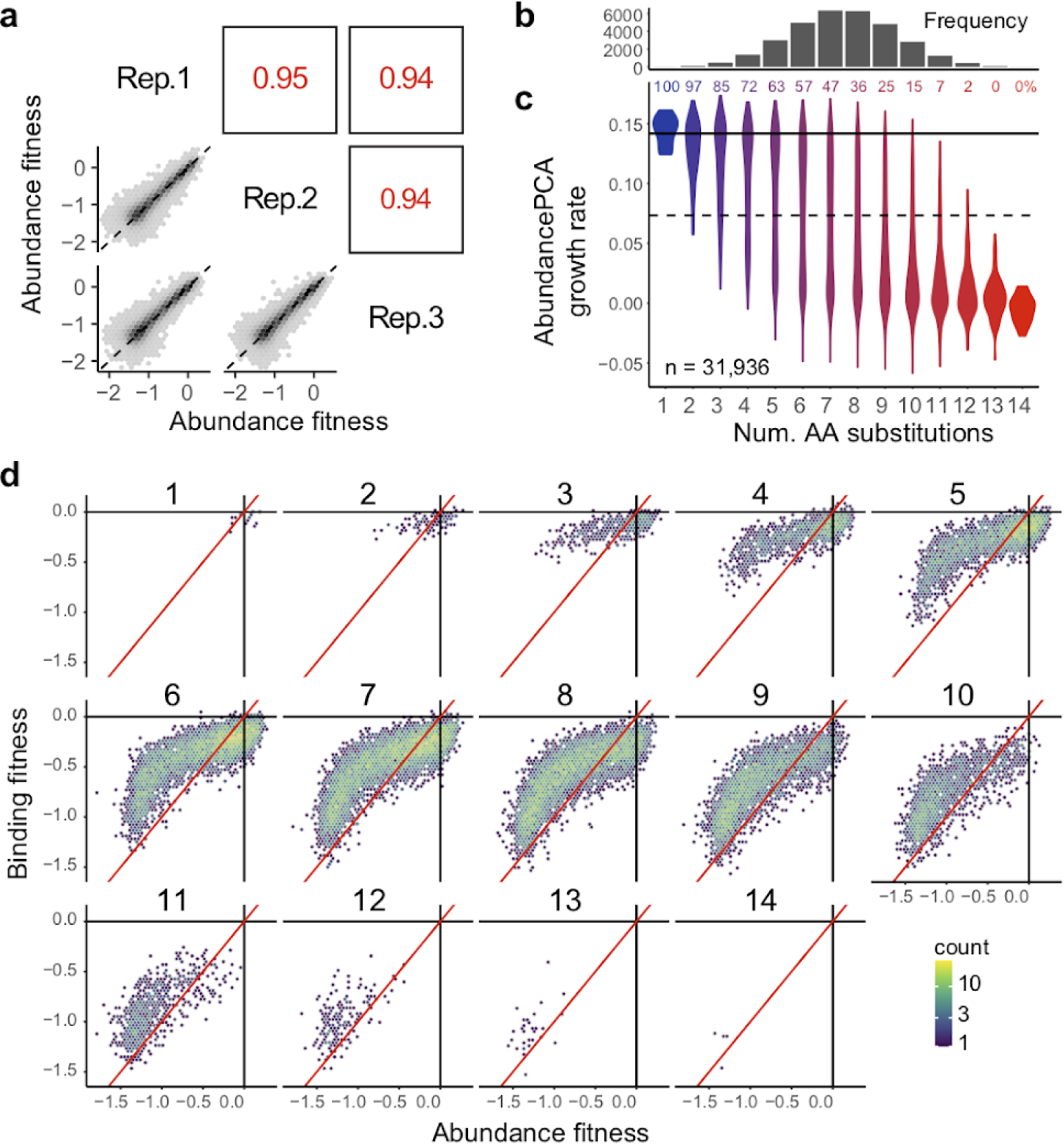
ddPCA data from combinatorial library 3 reveals that abundant multi-mutants are binding-competent (have conserved fold). **a**, Scatter plots showing the reproducibility of fitness estimates from AbundancePCA from combinatorial library 3 (see Fig. 5). Pearson’s r indicated in red. Rep., replicate. **b**, Histogram showing the number of observed aa variants at increasing Hamming distances from the wild type where the x-axis is shared with panel c. **c**, Violin plot showing distributions of AbundancePCA growth rates inferred from deep sequencing data versus number of amino acid substitutions. The percentage of bound protein variants (predicted fraction bound molecules > 0.5) is shown at each Hamming distance from the wild type. **f**, 2d density plots comparing abundance and binding fitness for increasing Hamming distances 1-14 from the wild type as indicated.

